# Using spatial statistics to infer game-theoretic interactions in an agent-based model of cancer cells

**DOI:** 10.1101/2025.07.09.664005

**Authors:** Sydney Leither, Maximilian A. R. Strobl, Jacob G. Scott, Emily Dolson

## Abstract

Drug resistance in cancer is shaped not only by evolutionary processes but also by eco-evolutionary interactions between tumor subpopulations. These interactions can support the persistence of resistant cells even in the absence of treatment, undermining standard aggressive therapies and motivating drug holiday-based approaches that leverage ecological dynamics. A key challenge in implementing such strategies is efficiently identifying interaction between drug-sensitive and drug-resistant subpopulations. Evolutionary game theory provides a framework for characterizing these interactions. We investigate whether spatial patterns in single time-point images of cell populations can reveal the underlying game theoretic interactions between sensitive and resistant cells. To achieve this goal, we develop an agent-based model in which cell reproduction is governed by local game-theoretic interactions. We compute a suite of spatial statistics on single time-point images from the agent-based model under a range of games being played between cells. We quantify the informativeness of each spatial statistic and demonstrate that a simple machine learning model can classify the type of game being played. Our findings suggest that spatial structure contains sufficient information to infer ecological interactions. This work represents a step toward clinically viable tools for identifying cell-cell interactions in tumors, supporting the development of ecologically informed cancer therapies.

**Author summary:** Drug resistance is a major challenge in cancer treatment, often leading to relapse despite initially successful therapy. While mutations are a key driver, ecological interactions between drug-sensitive and drug-resistant cells also play a critical role. These interactions are complex and dynamic, and few molecular biomarkers exist, making them difficult to study and account for in treatment planning. We use evolutionary game theory, a framework for quantifying interactions between cells, to investigate whether it is possible to infer these interactions using just a single time-point image of the cells. We develop an agent-based model where cells reproduce based on local interactions and quantify the resulting patterns in how cells are distributed across space using a suite of spatial statistics. We find that specific interaction types produce distinct spatial patterns that are evident in these metrics, and we train a simple machine learning model to classify the interaction type based on the metrics. Our results suggest that spatial data alone can offer valuable insights into tumor dynamics, potentially enabling more informed and adaptable cancer treatments based on eco-evolutionary principles.

## 1 Introduction

Cancers are heterogeneous and evolving diseases. Genetic and epi-genetic mutations give rise to clonally related, but geno- and phenotypically distinct subpopulations, which fuel the evolutionary processes by which cancers escape the body’s control and too often overcome even our most advanced treatments. Great progress has been made in understanding the mutational processes by which new clonal populations arise, and in molecularly characterizing these populations. However, cancer cells do not grow in isolation - they interact with each other, and other cells in their microenvironment, and there is growing evidence that these ecological interactions are crucial for understanding tumor evolution.

For example, Marusyk et al. [1] identified an IL11 expressing cancer subpopulation in a xenograft model of breast cancer which was vital for supporting growth of the tumor as a whole. Even though this subpopulation made up only 5.5% of the total population, eliminating this subpopulation caused tumor expansion to seize. Similar observations about one population supporting the growth of another were also made by others [2, 3, 4, 5]. Maltas et al. [5] used mathematical modeling to further demonstrate that such positive interactions significantly increase pre-treatment heterogeneity and thereby the probability of pre-existing, drug-resistant mutants, which in themselves may be selected for during tumor evolution. Furthermore, others have shown that targeting or leveraging these interaction holds new therapeutic opportunities. Studies in colorectal [6], lung [7], and ovarian [8] cancer have shown that drug-sensitive cells can suppress the outgrowth of drug-resistant cells, which has motivated so-called “adaptive therapy” (AT), which seeks to prevent or delay the fixation of the resistant population by taking eco-evolutionary dynamics into account via treatment holidays [9]. While the results of a pilot clinical trial in prostate cancer are promising [10], *in vitro* experiments in other cancers have shown that the extent to which resistant cells can be competitively suppressed depends on spatial constraints [6], nutrient constraints [8], and drug concentration [11]. Furthermore, sometimes resistant cancer cells out-compete sensitive cells regardless of drug [12]. Together, these examples show that ecological interactions play an important role in deciding which clonal populations survive and expand, and that deeper knowledge of these interactions holds promising therapeutic potential.

We need tools to measure ecological interactions in cancer and predict the resulting ecological dynamics. In ecology and evolution research, evolutionary game theory (EGT) is a powerful theoretical framework for understanding and predicting population dynamics [13]. There has been growing interest in applying it to cancer evolution [14, 15]. In EGT, ecological interactions are represented through the assumption that a population’s fitness is *frequency-dependent*, i.e. it depends not just on its intrinsic fitness but also on the frequency of other populations in the environment [13, 15, 16]. By assuming specific functional forms for this relationship, EGT allows predictions about the long-term evolutionary outcome - the evolutionary stable state - and its dependence on different model parameters (e.g. the strength and direction of interactions; [14]). For example, prior work has used EGT to model the role of cooperation [17, 18] and space [19, 20] in cancer evolution, and EGT modeling informed the design of the pilot trial of AT in prostate cancer [10, 21].

In this study we will focus on so-called symmetric, linear, two-strategy games [22], which provide a simple and powerful framework for understanding ecological interactions between two cell populations with distinct (heritable) phenotypes (referred to as “strategies” in game theory literature). While our results can be applied to any pair of cell populations, we are particularly interested in the role of ecology in drug resistance evolution and will therefore consider a drug-sensitive and drug-resistant tumor population. Linear games assume that the relationship between frequency and fitness is linear, and we can thus classify interactions as positive, negative, or neutral depending on whether the presence of one population increases, decreases, or does not affect the fitness of the other population, respectively (e.g. sensitive cells increasing, decreasing, or not affecting the fitness of resistant cells; Figure 1a).

**Fig 1.**
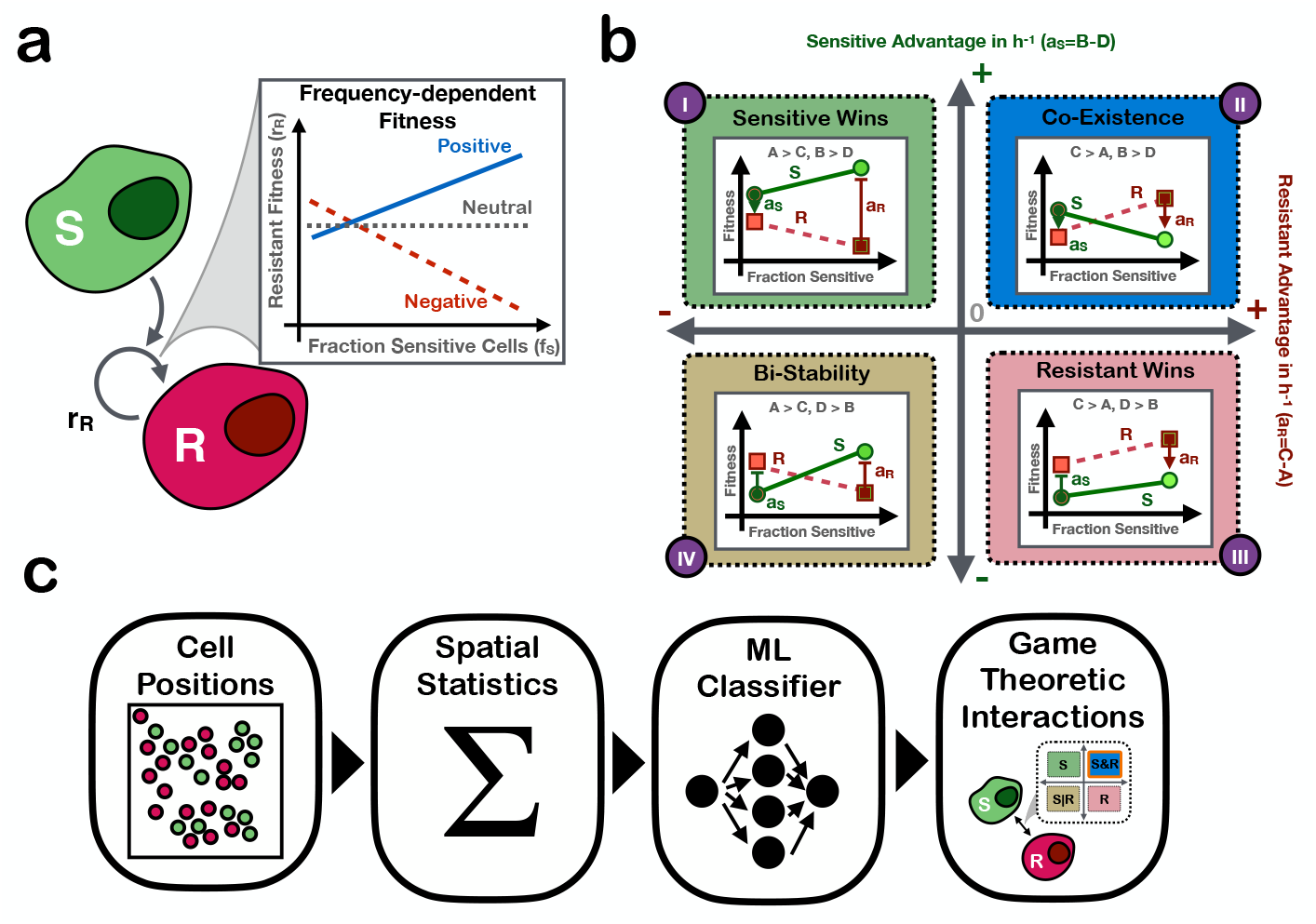
A visualize summary of the paper. **a)** ecological interactions between subpopulations may affect fitness; here we see the increased presence of sensitive cells may cause the resistant fitness to increase (positive interaction), decrease (negative interaction), or not change (neutral interaction). **b)** the game space and an example behavior expected in each quadrant. Behaviors are represented in terms of the growth rate for sensitive (green, solid) and resistant (pink, dashed) cells across population compositions. **c)** the pipeline presented in this paper: starting with a 2D single timepoint image of drug-sensitive and drug-resistant cells, we use spatial statistics from the image to infer the game space the cells’ interactions reside in.

To leverage EGT for real-world prediction-making, it is critical to develop methods to calibrate EGT models with empirical data. A common way to study cancer ecology is through co-culturing cell populations *in vitro* or *in vivo*, and measuring how each population’s growth rate in co-culture differs to their mono-culture growth rate. Several of the aforementioned studies have used this approach to classify interactions into positive, negative or neutral [2, 4, 12]. Extending this approach, Kaznatcheev et al [12] have recently developed the “game assay”, which uses a range of co-culture experiments starting from different initial seeding proportions to empirically measure the relationship between growth rate and population frequency. Using the game assay, they showed that linear games exist in cancer, and that the addition of drug or fibroblasts can alter how the game will evolve. Furthermore, Farrokhian et al [11] demonstrated that the concentration of drug can also alter the game being played by the cells, suggesting that through careful adjustment of drug levels it might be possible to steer tumor evolution. While co-culture experiments are powerful for directly establishing the impact of ecology on cell fitness, and investigating the underlying molecular mechanisms, they have several drawbacks: 1) the necessary experiments are resource-intensive, 2) the need to culture cells makes this technique challenging *in vivo* and in a clinical context, and 3) because cells evolve over time, the interactions between the cells may be evolving over time as well, making time series data complex to interpret.

To address these problems, this study investigates how much spatial patterns from single time-point imaging data reveal about the interaction between drug-sensitive and drug-resistant cells (Figure 1c). We develop an agent-based model whose agents (cells) reproduce based on the game played between themselves and each other cell in their local neighborhood to determine if we can use spatial statistics to infer the game the cells were playing based on a single time-point image. Since there is not currently a method to measure the true game real-world cells are playing, an agent-based model is necessary in order to generate data with ground-truth games to evaluate the game inference method. We analyze the aggregated spatial statistics present across samples from a diverse suite of payoff matrices and quantify how much information each spatial statistic reveals about the underlying game being played between the cells. We then utilize a simple machine learning model to infer the game based on the spatial statistics. As far as we are aware, this work represents the first excursion into identifying games based on single time-point spatial data.

## 2 Methods

### 2.1 Software and data availability

The agent-based model, analysis code, and instructions for reproducing the results are publicly available on GitHub ZENODO.

### 2.2 Game space

Thanks to the assumption of linearity, the effect of different cells interacting with each other can be represented by a payoff matrix that specifies the outcomes (in terms of growth rate) of interactions between all possible pairs of phenotypes as follows:

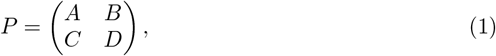

where a drug-sensitive cell has a payoff of *A* when interacting with another sensitive cell and payoff of *B* when playing against a resistant cell; *D* and *C* are the corresponding payoffs for a resistant cell. This means that when sensitive cells are primarily surrounded by other sensitive cells, they will grow at rate *A*, whereas when they are primarily surrounded by resistant cells they will grow at rate *B*, etc. Because most cells interact with more than one other cell, applying EGT to a population generally requires averaging payoffs across a set of interactions. In some cases, this calculation may assume the game is played with every other cell in the entire population, while in others it may be assumed to be played in only a local neighborhood [23]. Using this formulation, there are four possible long-term outcomes which are derived based on the payoff matrix [11].

We group the different games the cells could play into a “game space”, where the x-axis represents the “advantage” of resistant cells, *a*_*R*_, when invading into a population of sensitive cells (i.e. when they are primarily surrounded by sensitive cells; *a*_*R*_), and vice versa for the y-axis (*a*_*S*_; Figure 1 b). If *a*_*R*_ *>* 0, then a single resistant cell placed into a population of sensitive cells will expand and establish a population of resistant cells, whereas if *a*_*R*_ *<* 0 it will go extinct, and correspondingly for *a*_*S*_. In this way, games can be classified into one of four quadrants in the game space, shown in Figure 1 b): i) Q1 (*Sensitive wins*), in which *a*_*S*_ > 0 and *a*_*R*_ < 0, so that sensitive cells will always out-compete resistant cells, ii) Q2 (*Coexistence*), in which both sensitive and resistant cells grow faster playing against their opposite type than playing against their same type (*a*_*S*_, *a*_*R*_ *>* 0), resulting in stable coexistence of both types, iii) Q3 (*Resistant Wins*), in which resistant cells always grow faster than sensitive cells (*a*_*S*_ *<* 0, *a*_*R*_ *>* 0), and iv) Q4 (*Bistability*) where both cell types grow fastest around their same type (*a*_*S*_, *a*_*R*_ *<* 0), so that one type will competitively exclude the other depending on the initial population fraction (unless cells are seeded exactly at the unstable mixed equilibrium).

### 2.3 Agent-based model

In order to analyze spatial patterns of cells where the game-theoretic cell-cell interactions are known, we use an agent-based model (ABM). We create a 2D ABM in Java 1.8 using the Hybrid Automata Library [24], based on the ABM in [25]. We use a 100×100 lattice grid with no-flux boundary conditions. The agents in the ABM occupy a single site and represent the cells composing a tumor. We have two types of cells, which we interpret as drug-sensitive and drug-resistant. We define a cell’s local neighborhood as the set of cells within a Manhattan distance of two from it. Cells reproduce into one of four sites adjacent to it. The probability that a cell will reproduce at each time step is determined by the average payoff of the games played between a focal cell and the cells in its local neighborhood. If the four sites immediately adjacent to a cell are full, it cannot reproduce. Cells cannot move. Each cell has a 0.009 probability of death at each time step [26]. The game quadrants and an example of how cells reproduce based on their local neighborhood is shown in Figure 1b.

### 2.4 Agent-based model data acquisition

In order to extract comprehensive spatial patterns that could be used to infer the game that is defining cells’ interactions, we collected spatial data from multiple runs of the ABM. We ran 2500 samples of the ABM across different values of the payoff matrix parameters *A, B, C, D*, initial density, and initial proportion resistant. The payoff matrix parameters ranged from 0 to 0.1, rounded to two decimal places. To ensure both cell types were present, initial density ranged from 5% to 95% and initial proportion resistant ranged from 0.2 to 0.8. We used latin hypercube sampling [27] to evenly sample from this parameter space. Spatial data was collected after 200 timesteps. Timesteps could be interpreted as hours, as the range of the game space coordinates was chosen to correspond with the range of coordinates from Kaznatcheev et al [12], where the growth rate was measured in hours. Any samples where the game was undefined, because *A* equaled *C* or *B* equaled *D*, were discarded, leaving 2048 samples with defined games.

### 2.5 Spatial statistics

We extracted a set of 12 spatial statistics from the ABM spatial data and evaluated the spatial statistics’ performance in distinguishing games from each other. We use a range of existing spatial statistics, the full list of which can be found in Supplementary Table S1. These spatial statistics are extracted using their implementations in the MuSpAn package [28]. We also introduce a novel spatial statistic, the neighborhood composition. The description of this spatial statistic can be found in Supplementary Section S1.

Some spatial statistics used here are distributions or functions, rather than single values (e.g. the cross-pair correlation function). We convert these measurements into summary statistics to make analysis more interpretable and to enable them to be used as features in a machine learning model. Distribution spatial statistics are summarized as their mean, standard deviation, skew, and kurtosis. Function spatial statistics are summarized as their minimum and maximum [29]. The summary statistics of the distribution and function spatial statistics, along with the single-value spatial statistics, are throughout the manuscript referred to as features. We only retain one feature from each set of correlated features (Spearman’s rank correlation coefficient *ρ >* 0.9 [30]), for a total of 26 non-correlated features. The correlated feature clusters can be found in Supplementary Section S2. Features are z-score normalized to have unit variance.

## 3 Results

Interactions between drug-sensitive and drug-resistant cells play a crucial role in the emergence and maintenance of drug resistance in cancer. In order to incorporate the effect of interactions into cancer treatment, there must be tools that can effectively measure the interactions between subpopulations. Here, we present an analysis into using spatial statistics extracted from a single-timepoint image to infer the type of interaction between two cell types. We consider four types of interactions: *Sensitive Wins, Coexistence, Resistant Wins*, and *Bistability*, which we refer to as the game, motivated by evolutionary game theory (Figure 1). We use an agent-based model to generate 2048 images of cells with ground-truth games. Cells in the agent-based model express the game by reproducing with probabilities determined by the cell composition of their local neighborhood and the defined payoff matrix (Equation 1).

### 3.1 Spatial patterns are qualitatively different across agent-based games

As is evident from figure 2, different games cause the ABM (Section 2.3) to produce qualitatively distinct spatial patterns of sensitive and resistant cells. As expected, in *Sensitive Wins* and *Resistant Wins*, the respective cell types approach fixation. Because cells in *Coexistence* grow fastest when near the opposite cell type, these games lead to the cell types being highly intermixed. In *Bistability* the cells exhibit more clumping than any other game quadrant, since cells grow fastest when near other cells of the same type. This example illustrates that different games generate distinct spatial patterns.

**Fig 2.**
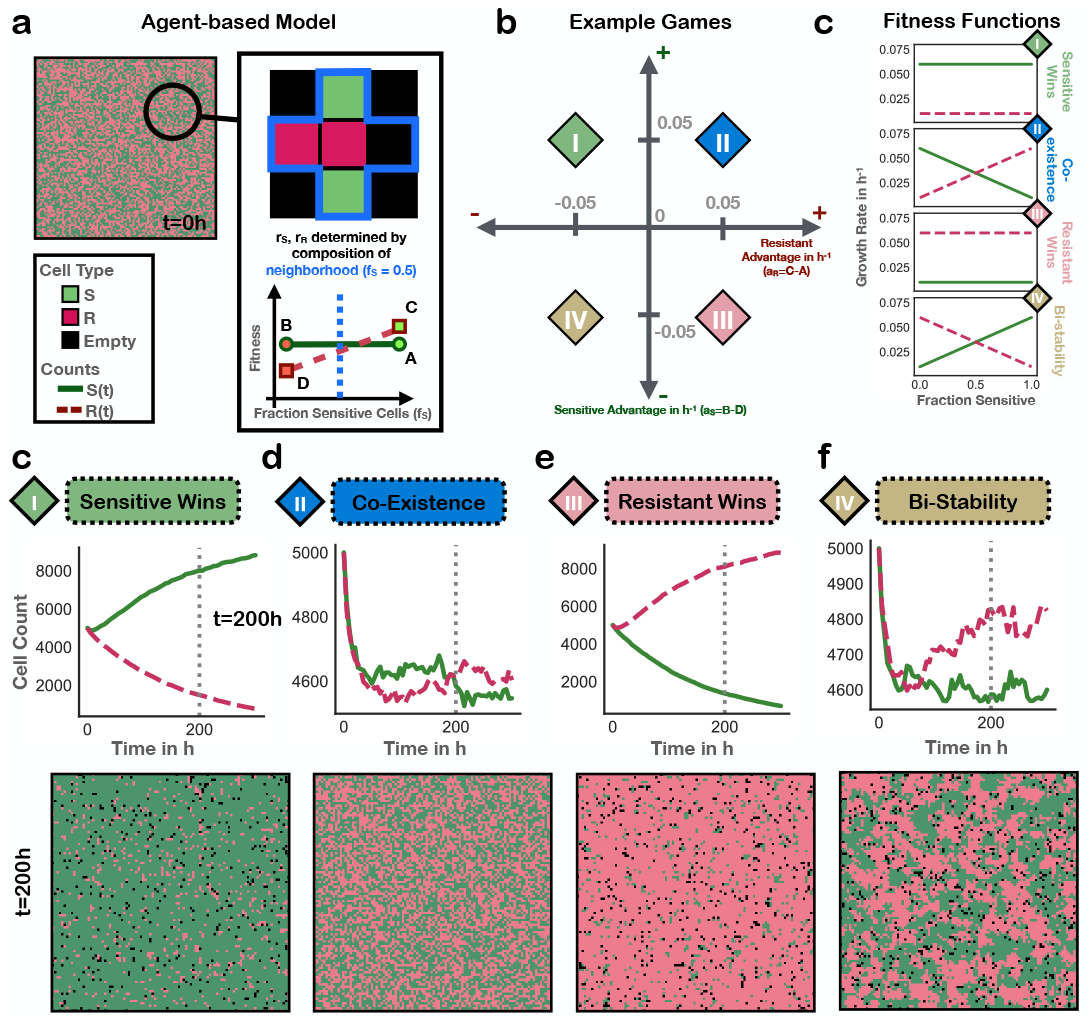
Visual emergence of games from the same initial conditions. **a)** A visual example of how cells reproduce in the ABM. Resistant and sensitive cells are colored pink and green, respectively. **b)** The game space and **c)** example fitness functions for each quadrant. **d)** The image in a) is the state of the ABM at the start of the runs, with 100% initial density and 50% fraction resistant. The bottom of d) shows ABM states after 200 timesteps when running the same starting conditions but changing which game quadrant the payoff matrix resides in. The payoff matrix parameters were set such that each value was either 0.06 or 0.01, in a combination which would place each payoff matrix into a unique game quadrant.

### 3.2 Pairs of games are distinguished by different spatial statistics

As part of our goal to distinguish game based on spatial statistics, we investigate which spatial statistics provide the most distinguishability between pairs of games and why. To systematically quantify differences in the spatial patterns between games, we sampled 2048 different games and in each case collected a set of 12 spatial statistics from a snapshot of the ABM after 200 timesteps. Games were sampled by varying the location in the game space, initial density, and initial fraction sensitive using latin hypercube sampling (Section 2.4). Spatial statistics are converted into single-valued features using summary statistics (Section 2.5). To identify which features best indicate the type of game the cells are playing (e.g. *Sensitive Wins*), we compile the distribution of each feature across samples within each game. We then analyze the distinguishability of the feature distributions for each pair of games. We calculate distinguishability using the Earth movers distance (EMD) [31], also called the Wasserstein distance, which measures the dissimilarity between two distributions. Figure 3 shows the three features with the highest EMD for each pair of games, colored by overall spatial statistic (for the full list of features’ EMDs for each pair of games, see Supplementary Section S3). We see that each pair of games have different features that distinguish them the best. Some features require a focal cell type and are asymmetric, meaning that the relationship between sensitive and resistant is different from the relationship between resistant and sensitive. In these cases, we denote the focal type with labels in the format “SR,” which means that the sensitive cell (S) was the focal cell type, while the resistant cell (R) was the non-focal. Features that only act on one cell type are labeled with that full cell type, i.e. “sensitive” or “resistant”. We now consider some specific spatial statistics that proved to be important.

**Fig 3.**
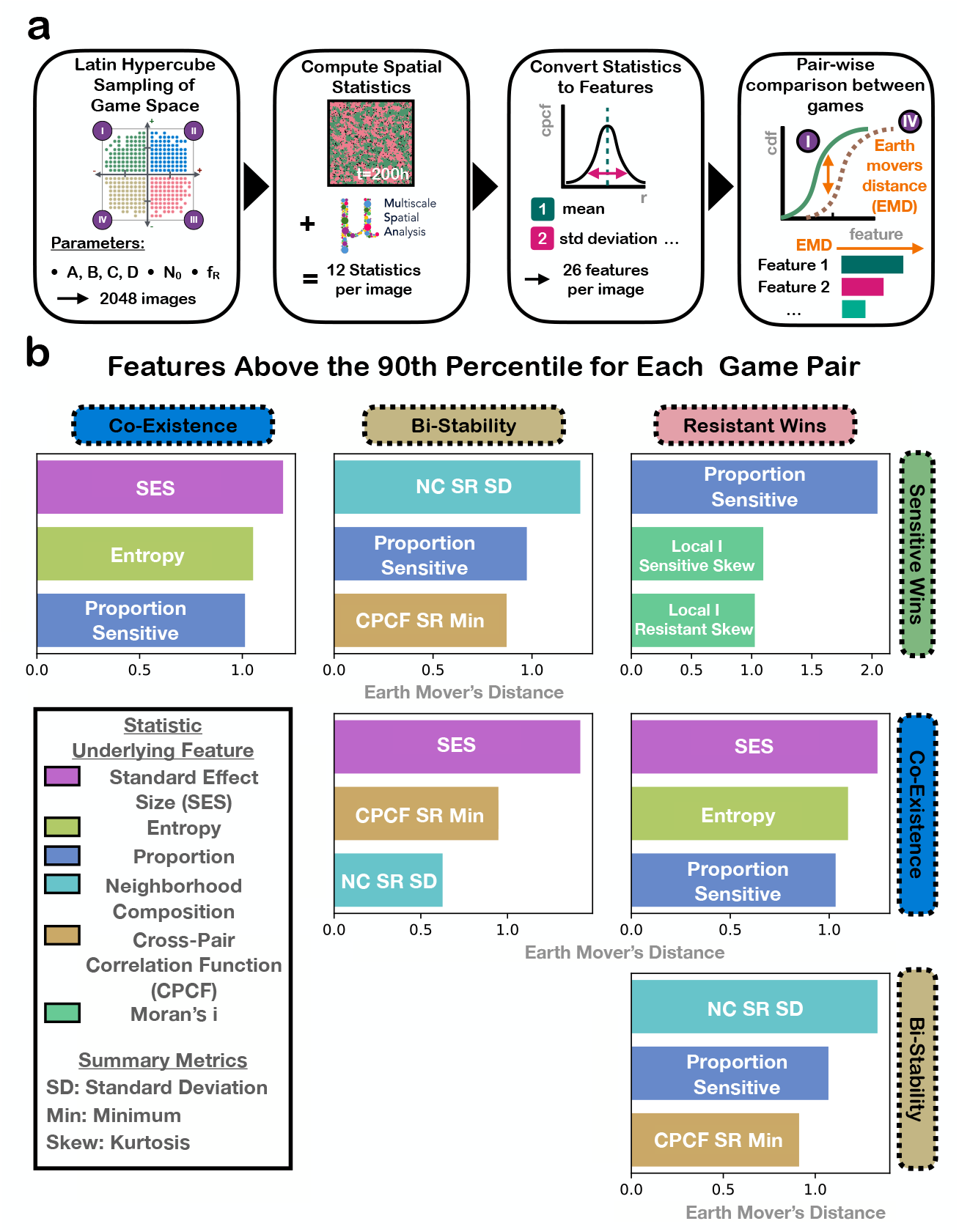
Distinguishing spatial statistic features above the 90th percentile for each pair of games. a) A visual explanation of how b) was created. **b)** Distinguishing spatial statistic features above the 90th percentile for each pair of games. Distinguishability is calculated as the Wasserstein distance between feature distributions. Features are colored by overall spatial statistic type. “SR” means sensitive was the focal cell type in the spatial statistic calculation, vice versa for “RS”.

#### Proportion sensitive

Proportion sensitive is seen throughout Figure 3 as one of the top distinguishing features for multiple game pairs. Proportion sensitive, which detects the proportion of sensitive cells in the population, distinguishes the best between *Sensitive Wins* and *Resistant Wins* (Figure 3). Since these games promote fixation of their respective cells, proportion sensitive can efficiently capture the imbalance in cell types that suggests *Sensitive Wins* or *Resistant Wins. Coexistence* tends to have more intermediate values of proportion sensitive, as each cell type grows faster around the opposite cell type, stably maintaining both cell types in the population (Supplementary Figure S7). This mutualistic growth leading to a more equal proportion of cell types explains why proportion sensitive also distinguishes both *Resistant Wins* and *Sensitive Wins* from *Coexistence. Bistability* has the widest range of proportion sensitive values seen in the samples, as cells grow fastest around their same type, encouraging local competition between the types that may lead to the extinction of the less populous type (Supplementary Figure S7). Thus proportion sensitive is not useful for distinguishing *Bistability* from the other games.

#### Local Moran’s i Skew

Local i Sensitive Skew and Local i Resistant Skew, refer to the the skew of the Local Moran’s i statistics of the sensitive cell and resistant cell respectively, and are top features for distinguishing between *Sensitive Wins* and *Resistant Wins*. Local Moran’s i is calculated by overlaying a hexagonal grid on the cell layout, counting the focal cell types within each hexagon, and computing the local Moran’s i statistic to assess similarity with neighboring hexagons. A positive Moran’s i value indicates clustering of the focal cell type, while a negative Moran’s i value indicates dispersion. The skew of the distribution of all local Moran’s i values indicates how biased the data is to exhibit more clustered areas than dispersed or vice versa. In Supplementary Figure S7, we see that the distribution of Local i Sensitive Skew for *Resistant Wins* tends to have larger values than the other distributions, and vice versa for Local i Resistant Skew for *Sensitive Wins*. We hypothesize that this difference reflects the fact that the non-focal cell type tends to survive for longer when in monotypic clusters.

#### Neighborhood Composition Standard Deviation

The neighborhood composition (NC) metric quantifies the average cell composition in the local neighborhood of a given cell type. The feature NC SD is the standard deviation of this metric across all instances of the same game. NC SD summarizes the diversity of neighborhood compositions of the focal cell type. NC RS SD, the standard deviation of the neighborhood composition when the resistant cell is the focal cell type, distinguishes *Bistability* from *Resistant Wins*, and NC SR SD distinguishes *Bistability* from *Sensitive Wins* (Supplementary Figure S7). *Bistability* tends to have diverse neighborhoods as local regions of the grid tend towards either majority sensitive or resistant depending on the starting composition, which leads to a wide variety of observed neighborhood compositions. *Sensitive Wins* tends to have less diversity in sensitive cell neighborhoods, as most of the sensitive cells have a small number of resistant cells in their neighborhood. Thus, NC SR SD distinguishes *Bistability* and *Sensitive Wins*. The same logic applies for NC RS SD’s distinguishability of *Bistability* and *Resistant Wins*.

#### Cross Pair Correlation Function Minimum

The cross pair correlation function (CPCF) appears as one of the top three distinguishing features for all pairs of games including *Bistability*. CPCF compares the number of cells of the non-focal type present in an annulus of a given radius from each focal type cell to the number expected under a null hypothesis of complete spatial randomness. The CPCF is a measure of clustering or exclusion of the two cell types. *Bistability* tends to have a smaller CPCF Min than the other games (Supplementary Figure S7), indicating that cells of one type exclude cells of the other, as expected under *Bistability*. We hypothesize this is due to the fact that cells reproduce in their local neighborhoods, so that there is always a degree of exclusion between the cell types that may make it difficult for CPCF to distinguish *Coexistence* from *Resistant Wins* and *Sensitive Wins*.

#### Standard Effect Size of the Quadrant Correlation Matrix

Standard effect size of the quadrant correlation matrix (SES) has the highest distinguishability for *Coexistence* and each other game. The quadrant correlation matrix splits the image into hexagons and calculates how often the two cell types are seen together compared to what would be expected under complete spatial randomness. This feature captures that cell are more intermingled in *Coexistence* than in the other games (Supplementary Figure S7). SES may not distinguish *Bistability* from *Resistant Wins* and *Sensitive Wins* as these games often exhibit similar patterns of spatial segregation, where one cell type dominates local regions, reducing the frequency of co-localization that SES is designed to detect.

#### Entropy

Entropy, which we use to quantify the heterogeneity in the spatial arrangement of cell types, is one of the top features for distinguishing *Coexistence* from both *Sensitive Wins* and *Resistant Wins*. Higher entropy indicates a more mixed or disordered pattern, while lower entropy reflects more segregation or uniformity.

*Coexistence* exhibits high entropy because cells of different types are intermingled throughout the image, resulting in more disordered and heterogeneous spatial patterns (recall Figure 2). In contrast, both *Sensitive Wins* and *Resistant Wins* tend to show large regions dominated by a single cell type due to competitive exclusion, producing more ordered and predictable spatial patterns and therefore lower entropy (Figure 2). However, entropy does not distinguish *Coexistence* from *Bistability* (Supplementary Figure S2). In *Bistability*, cells cluster with their own type, forming large homogeneous patches (Figure 2). These patches vary in size and location across the image, creating moderate disorder and spatial heterogeneity. As a result, the entropy values of *Bistability* overlap with the other quadrants (Supplementary Figure S7).

### 3.3 Individually uninformative features can be synergistic

Since even the most distinctive feature distributions still overlap between games (as shown in Section 3.2), individual features alone don’t fully capture the spatial pattern differences. In this section, we use information theory [32, 33] to investigate whether pairing features can improve our ability to identify the game being played between cells. We start by examining the mutual information between each feature *F*_*i*_ and the game *G, I*(*F*_*i*_; *G*) = *H*(*F*_*i*_) −*H*(*F*_*i*_ | *G*), where *H* is the Shannon Entropy [32]. Mutual information quantifies the amount of information gained about the game by observing the given feature. In order to calculate the mutual information, which requires discrete variables, we discretize the features into equal-frequency bins. We calculate the number of bins for each feature using Sturges’ rule [34]: log *n* + 1, where *n* is the number of samples. We normalize the mutual information by the entropy of the game variable: 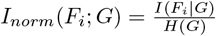.

Supplementary Figure S9 shows the mutual information between each feature and the game, colored by the spatial statistic the feature is derived from. Interestingly, the features with the highest mutual information are not always the features that best distinguished pairs of games (Figure 3). Thus, features that can differentiate well between pairs of games do not necessarily provide information that distinguishes among all games. The feature with the highest mutual information with the game is proportion sensitive at 0.29. This means that 29% of the information in the game can be explained by the proportion sensitive (since we normalized the mutual information by the game entropy). The fact that even the maximum mutual information from a single feature is so low suggests that a single feature cannot capture the spatial pattern differences between games.

To determine how much information a pair of features captures about the game, we can use interaction information. Interaction information [35, 36] measures the joint information of three variables. In our context, interaction information quantifies how much additional information about the game is provided by considering the interaction between two features. Interaction information is defined as:

*II*(*F*_*i*_, *F*_*j*_, *G*) = *I*(*F*_*i*_, *F*_*j*_; *G*) − (*I*(*F*_*i*_; *G*) + *I*(*F*_*j*_; *G*)) [36]. We also normalize the interaction information by the game entropy. Interaction information thus captures how much more is learned about a game when two features are considered together, beyond what is learned from each feature separately. A positive value indicates synergy, because the interaction between features provides information about the game that is not in either feature independently. A negative value indicates redundancy, because the features share the same information about the game. Figure 4 shows the interaction information for each pair of features. We see overall, Entropy, CPCF RS Min, and NC SR SD have the most synergy with other features. In the single feature analysis (Section 3.2), entropy was only effective at distinguishing *Coexistence* from *Sensitive Wins*/*Resistant Wins*. Similarly, both CPCF RS Min and NC SR SD were only effective at distinguishing *Bistability* from the other games. The high interaction information of these features suggests that the other features by themselves provide little information to distinguish between *Bistability* and *Coexistence*. We also see that proportion sensitive has redundancy with multiple features, meaning that those other features already contain information about proportion sensitive, and vice versa. 93% of the feature pairs exhibit synergy, which shows that pairs of features can compliment each other, providing explanatory power not possible with individual features. In the next step, we build on this observation by leveraging machine learning to develop higher-order combination of features with which to accurately and precisely identify the game from a single image.

**Fig 4.**
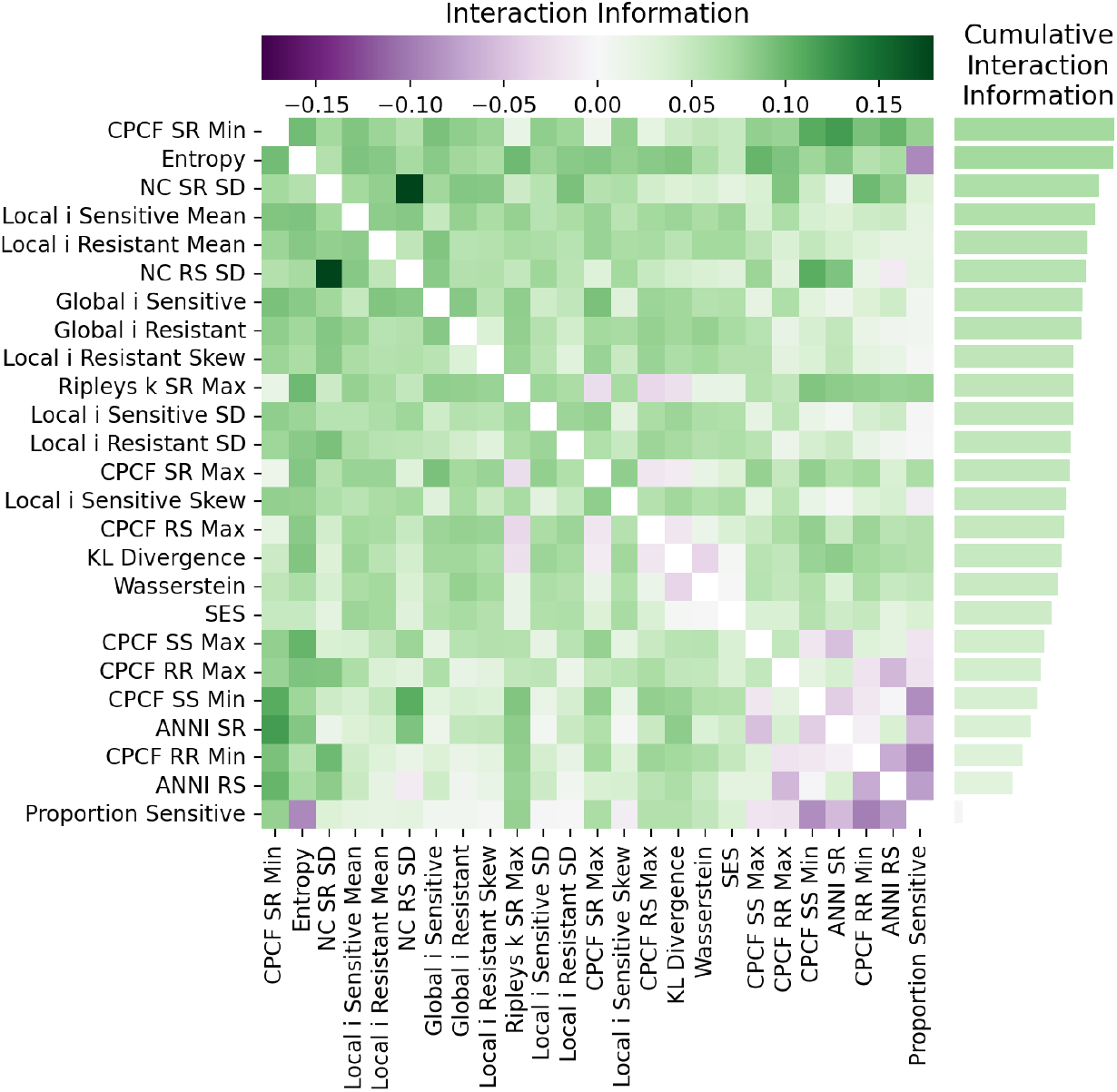
Interaction information between each pair of features and game. Interaction information between feature pairs quantifies how much additional information about game is gained by considering feature interactions beyond individual contributions. Positive values (synergy) indicate complementary information, while negative values (redundancy) suggest overlapping information.

### 3.4 Machine learning can infer agent-based game from spatial features

We use machine learning to analyze the ability of spatial features to predict the game being played between cells. We implement a simple neural network using Scikit-learn [37] with one layer, 300 nodes (Supplementary Figure S10), and default parameters from Scikit-learn. To identify the most informative features for the model, we employ a top-ten forward sequential feature selection procedure. This algorithm operates iteratively. In the initial step, a single-node, single-layer neural network is trained and evaluated (via 5-fold cross validation) using each feature individually. In the subsequent step, the ten top-performing features are each paired with another feature, and the model is re-evaluated for each resulting two-feature combination. The ten top performing combinations are retained. This process continues iteratively: at each stage, new feature sets are generated by appending one additional feature to each of the current top-performing sets, followed by model evaluation and selection of the best-performing combinations. The procedure terminates when no further improvement in model performance is observed among the top feature sets (Supplementary Figure S11).

Using this process, we find the best performing feature set to consist of 12 features (Supplementary Table S3). The average testing accuracy over 5-fold cross validation is 0.78. Supplementary Figure S12 shows the ROC curves. Figure 5 shows the confusion matrix of the testing data. We see *Coexistence* has the highest prediction accuracy, followed by *Resistant Wins, Sensitive Wins*, and *Bistability. Bistability* is commonly misclassified as *Sensitive Wins* or *Resistant Wins*, while *Sensitive Wins* and *Resistant Wins* are commonly misclassified as *Bistability. Sensitive Wins* and *Resistant Wins* are rarely misclassified as each other, and *Bistability* and *Coexistence* are rarely misclassified as each other. *Bistability* overall has the lowest classification accuracy, which makes intuitive sense given that *Bistability* exhibits unstable mixed equilibrium characteristics, i.e. multiple runs defined with the same payoff matrix may converge to either resistant or sensitive fixation depending on the starting conditions.

**Fig 5.**
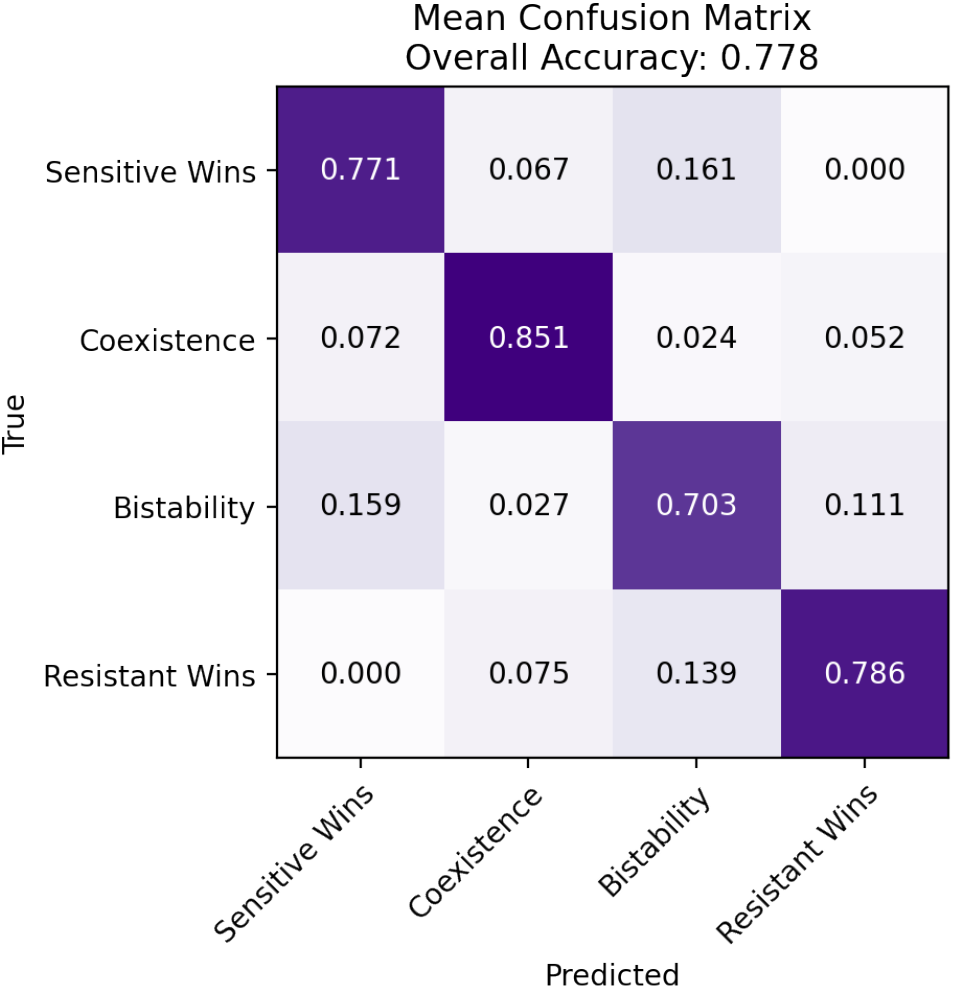
Mean testing accuracy confusion matrix over 5-fold cross-validation of a simple neural network trained on spatial statistic features. The model size and features included in the model were chosen as described in the results.

## 4 Discussion

Taken together, our results suggest that different game theoretic interactions are sufficient to produce quantifiably different spatial patterns. We find that different spatial statistics are important for distinguishing between each pair of game types. Furthermore, pairs of spatial statistics can contribute synergistic information with the game. Training a simple neural network model on a selected set of spatial statistic features reveals that, with the spatial statistics tested in this paper on our specific agent-based model data, *Coexistence* is the easiest game to classify, followed by *Resistant Wins*/*Sensitive Wins*, and then *Bistability*.

A critical question for future research is how well these results generalize from our ABM to *in vitro, in vivo*, and clinical settings. Theoretically, any system governed by spatial game theoretic interactions should exhibit similar general patterns to those observed here, but in practice there are a variety of reasons they might differ. In particular, although our Latin hypercube allowed us to explore a wide variety of model parameters, some real-world systems might have parameters outside the range of those we tested. For example, increasing the local neighborhood size and reproduction radius will push the system closer to a well-mixed state, weakening spatial patterns and the distinguishability of game by spatial statistics. Moreover, even among parameter sets that produce spatial patterns, the patterns may look different in different regions of the parameter space. For example, different cell lines may exhibit different growth behaviors (e.g. different rates of migration or spatial clustering), leading to different spatial patterns even across similarly conducted *in vitro* experiments. While the overall approach presented in this paper of analyzing pairwise game distinguishability, analyzing spatial statistic mutual information, and inferring games from spatial statistics should be valid across systems, the specific conclusions of which spatial statistics are informative, and how, may vary. To mitigate this gap between ABMs and the real world, the ABM could be modified with parameters and behaviors that better match the real-world cell behavior of specific cells. For example, to precisely and accurately model a particular cell type, it may be necessary to consider behaviors such as cells physically pushing each other, mutation, and diffusable growth factors. In the next step, we also plan to fit the ABM to real-world *in vitro* data for specific cell lines.

Game theory informed cancer treatment strategies are an obvious next step in evolutionary cancer therapy, an approach which has already made great strides [10]. The primary current iteration of these strategies, adaptive therapy, dynamically modulates treatment to preserve sensitive subpopulations [9]. However, this approach works best if, in the absence of drug, the cell types are playing a “Sensitive Wins” game, and could be detrimental if resistant cells are supported by sensitive cells. Measuring the game being played between the sensitive and resistant subpopulations, then, would give insight into whether this strategy is viable. Moreover, it opens the door to developing treatments that shift the game played to facilitate adaptive therapy. Yet so far, treating cancer based on game theory remains largely theoretical, in part due to the lack of practical tools for detecting game-theoretic interactions between cells. While the evolutionary game assay [11, 12] can measure payoff matrices between two subpopulations, it requires time-series data, limiting its clinical utility. Our work here suggests that it should be possible to instead infer the type of game being played between drug-sensitive and drug-resistant cells using only spatial information from a single time point. This approach is more compatible with clinical workflows, as it could eventually be run on tumor biopsy data. By making it possible to classify ecological interactions from a single time-point image, our method takes a step toward translating game-theoretic insights into real-world cancer treatment planning.

## Acknowledgments

Research reported in this publication was supported by the National Cancer Institute of the National Institutes of Health under award number U01CA280829. The content is solely the responsibility of the authors and does not necessarily represent the official views of the National Institutes of Health. This work was also supported in part through computational resources and services provided by the Institute for Cyber-Enabled Research at Michigan State University.

