## Supplement for "Using spatial statistics to infer game-theoretic interactions in an agent-based model of cancer cells"

### 1 Spatial statistics

#### 1.1 Neighborhood composition

The neighborhood composition (NC) spatial statistic quantifies the average neighborhood structure around each cell of a given type. NC is calculated as follows: for each cell of type A, record the fraction of type B cells in their local neighborhood. Remove any data points where the fraction of type B is zero. The resulting data set is the distribution of the fraction of cell type B in the neighborhood of each cell type A. In this work, the local neighborhood size for calculating NC was set to 3.

| Spatial Statistic | Type | Description | Source |
|---|---|---|---|
| Average Nearest Neighbor Index (ANNI) | Value | Clustering of cell types based on expected nearest neighbor distances | [1] |
| Cross Pair Correlation (CPCF) | Function | Aggregation of cell types across annuli | [2] |
| Cross Ripley's k | Function | Co-localization of cell types across radii | [3] |
| Entropy | Value | Shannon entropy of cell type A | [4] |
| KL Divergence | Value | Kullback-Leibler divergence between kernel density estimations of cell types | [5], [6] |
| Global Moran's i | Value | Spatial autocorrelation of continuous "proportion of cell type A in hex" over the whole population | [7] |
| Local Moran's i | Distribution | Spatial autocorrelation of continuous "proportion of cell type A in hex" in local neighborhoods | [7] |
| Nearest Neighbor (NN) | Distribution | Distances from each cell type A to any cell type B | [8] |
| Neighborhood Composition (NC) | Distribution | Fraction of cell type A in each cell type B's neighborhood | This paper |
| Proportion Sensitive | Value | Proportion of sensitive cells | This paper |
| Standard Effect Size of Quadrant Correlation Matrix (SES) | Value | Correlation between counts of cell types across regions | [9] |
| Wasserstein | Value | Wasserstein distance between 1D-projected spatial data of two cell types | [10] |

**Table 1. Spatial statistics used in this work.** Implementations of each spatial statistic not sourced from this paper are from MuSpAn [11]. Parameters of each spatial statistic were set to MuSpAn's default values divided by 10, as MuSpAn expects images around size 1000x1000 and our images were of size 100x100.

#### 2 Correlated feature clusters

| **Chosen Feature** | **Features in Correlated Cluster** |
|---|---|
| ANNI RS | |
| ANNI SR | |
| CPCF RR Min | |
| CPCF RR Max | |
| CPCF SR Min | CPCF RS Min, Ripleys k RS Min, Ripleys k SR Min |
| CPCF RS Max | |
| CPCF SR Max | |
| CPCF SS Min | |
| CPCF SS Max | |
| Entropy | |
| Global i Resistant | |
| Global i Sensitive | |
| KL Divergence | |
| Local i Resistant Mean | |
| Local i Resistant SD | |
| Local i Resistant Skew | |
| Local i Sensitive Mean | |
| Local i Sensitive SD | |
| Local i Sensitive Skew | |
| Proportion Sensitive | NC RS Mean, NC RS Skew, NC SR Mean, NC SR Skew, NN RS Mean, NN RS SD, NN RS Skew, NN SR Mean, NN SR SD, NN SR Skew, , Ripleys k RR Min, Ripleys k RR Max, Ripleys k SS Min, Ripleys k SS Max |
| NC RS SD | |
| NC SR SD | |
| Ripleys k RS Max | |
| Ripleys k SR Max | |
| SES | |
| Wasserstein | |

**Table 2. Correlated feature clusters.** Features are considered correlated if their Spearman's rank correlation coefficient $\rho > 0.9$. From each correlated cluster, we only analyze the chosen feature in this work.

#### 3 Pairwise game distributions

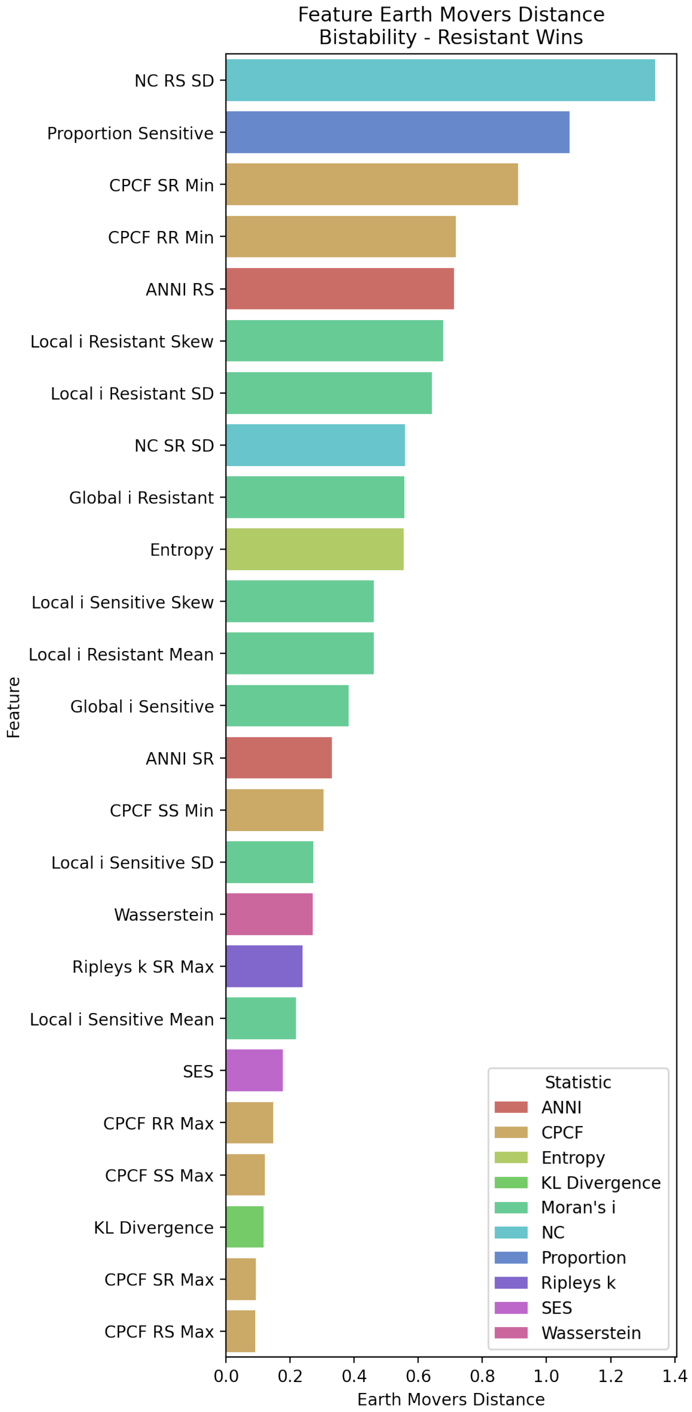


**Fig 1. Full list of the distinguishability (Earth movers distance) between the feature distributions of *Bistability* and *Resistant Wins*.**

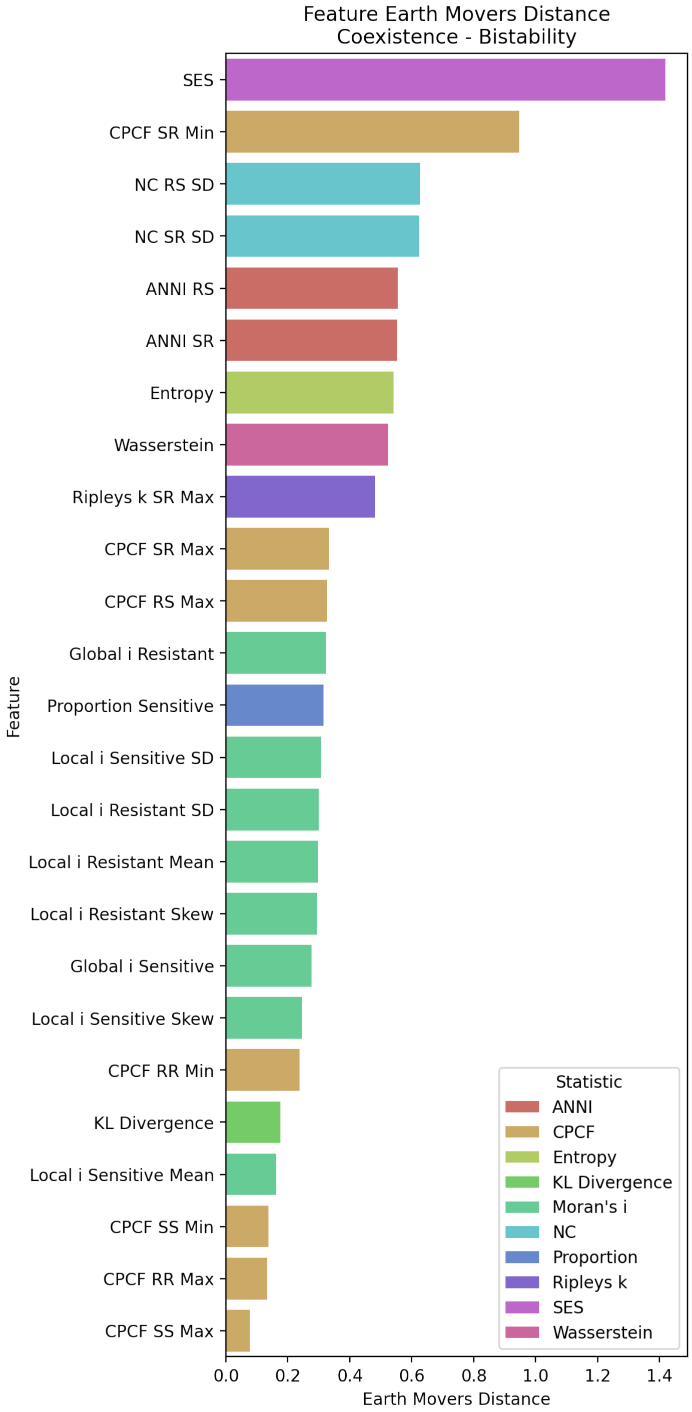


**Fig 2. Full list of the distinguishability (Earth movers distance) between the feature distributions of *Coexistence* and *Bistability*.**

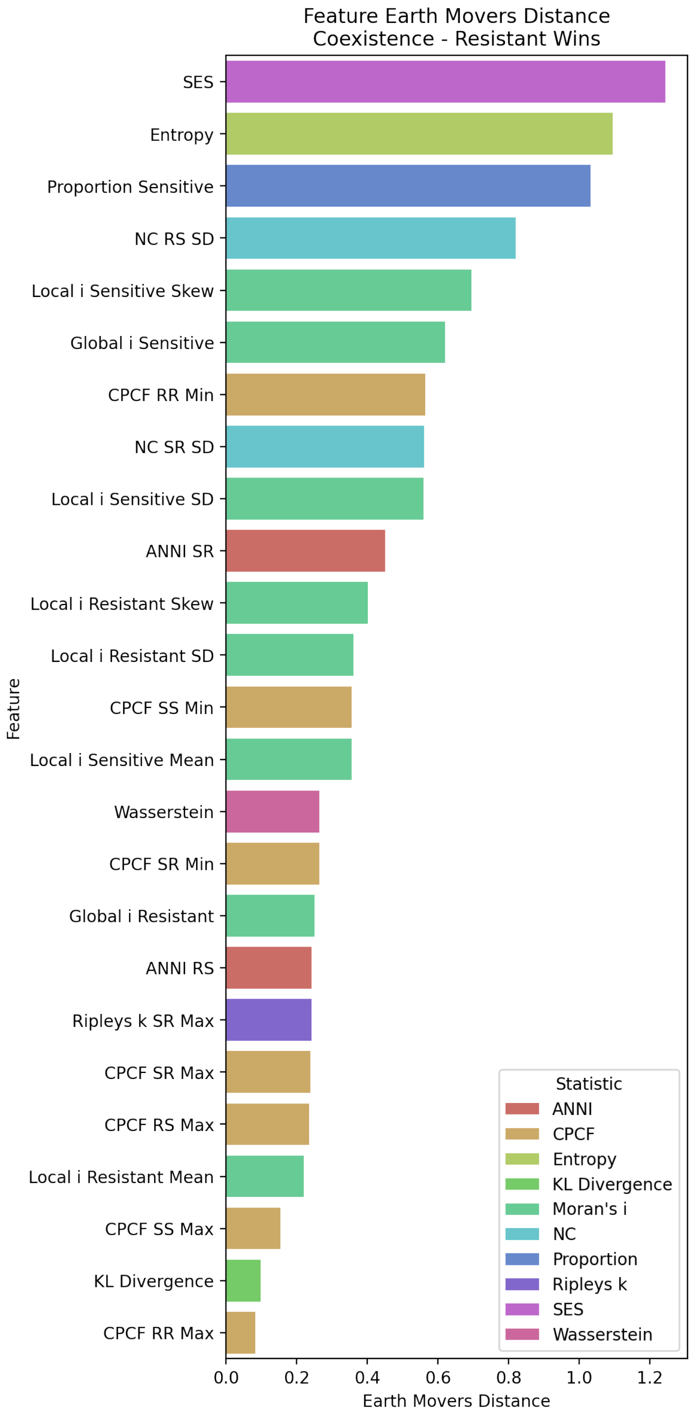


**Fig 3. Full list of the distinguishability (Earth movers distance) between the feature distributions of *Coexistence* and *Resistant Wins*.**

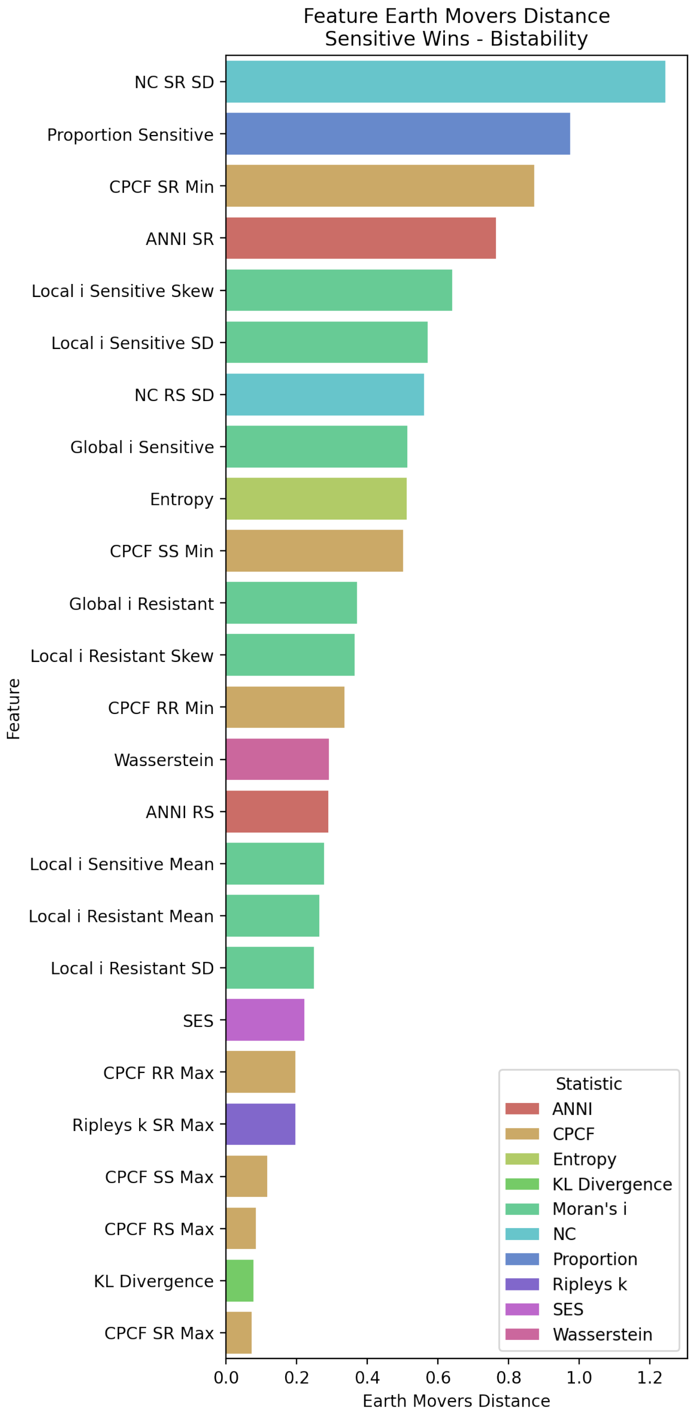


**Fig 4. Full list of the distinguishability (Earth movers distance) between the feature distributions of *Sensitive Wins* and *Bistability*.**

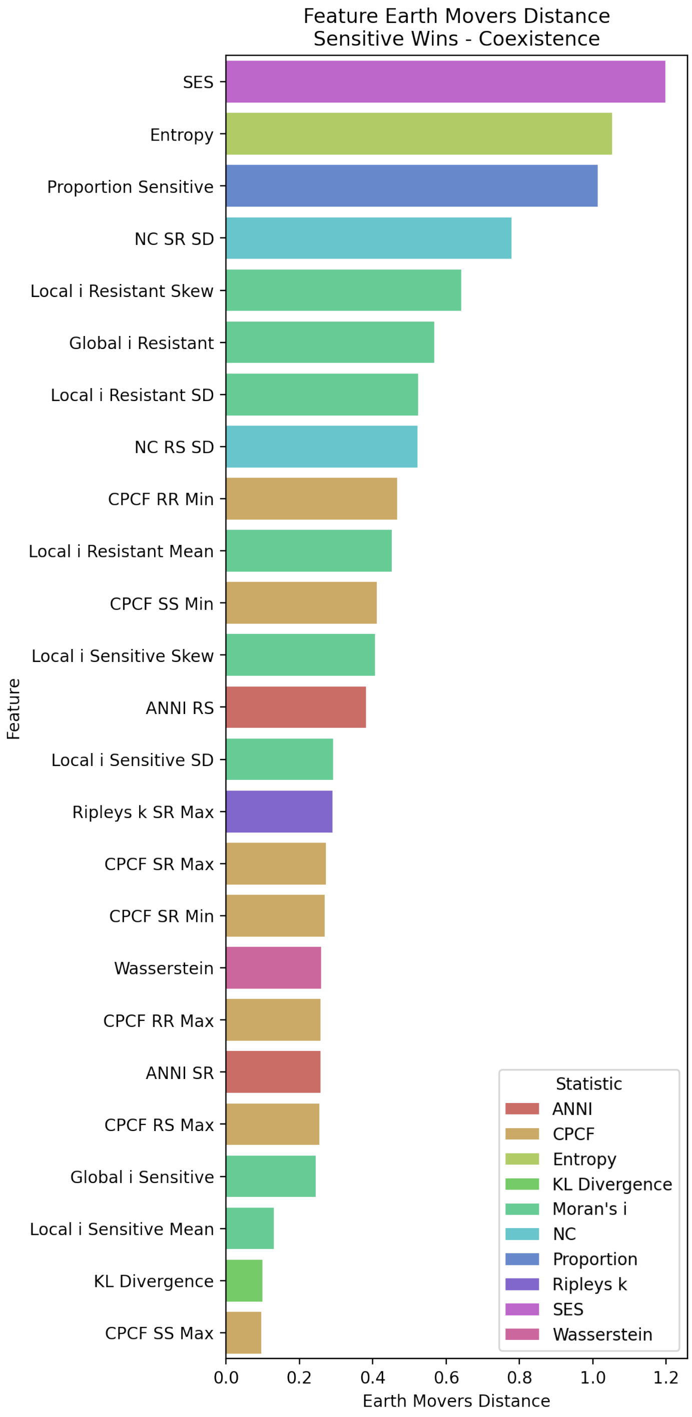


**Fig 5. Full list of the distinguishability (Earth movers distance) between the feature distributions of *Sensitive Wins* and *Coexistence*.**

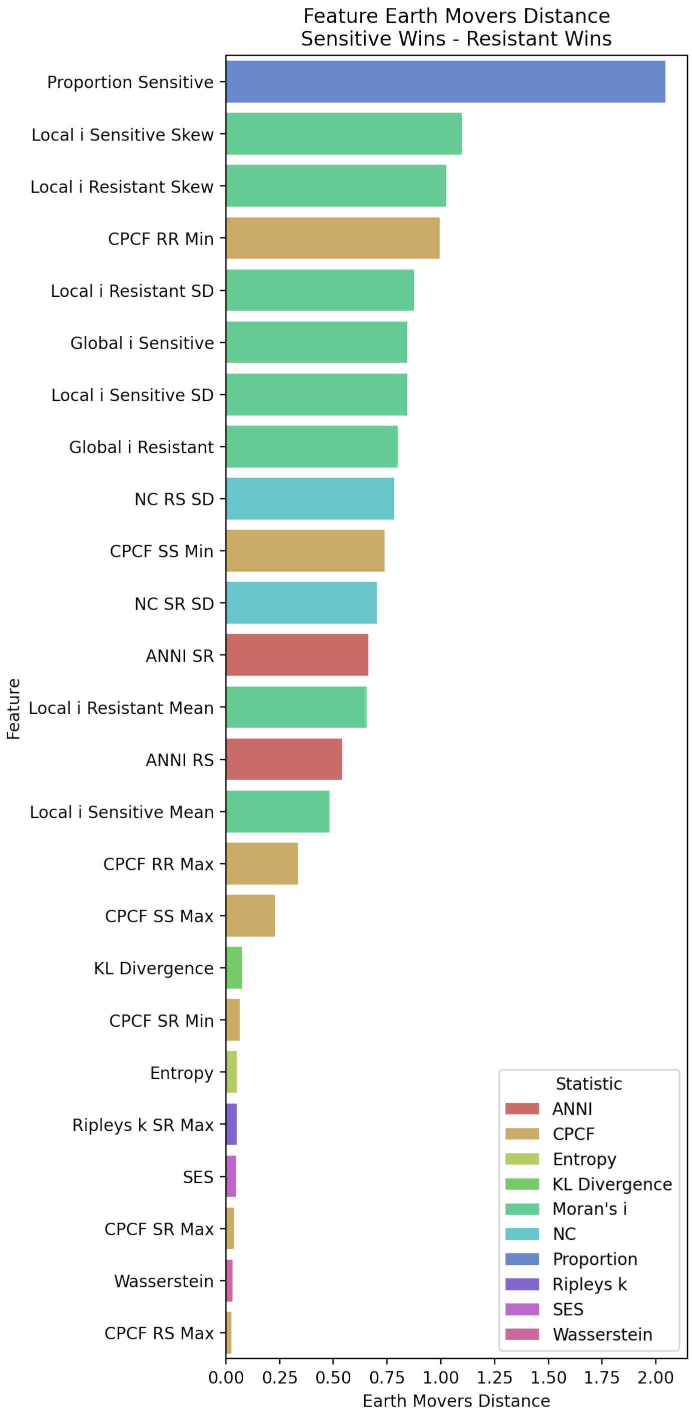


**Fig 6. Full list of the distinguishability (Earth movers distance) between the feature distributions of *Sensitive Wins* and *Resistant Wins*.**

#### 4 Feature distributions

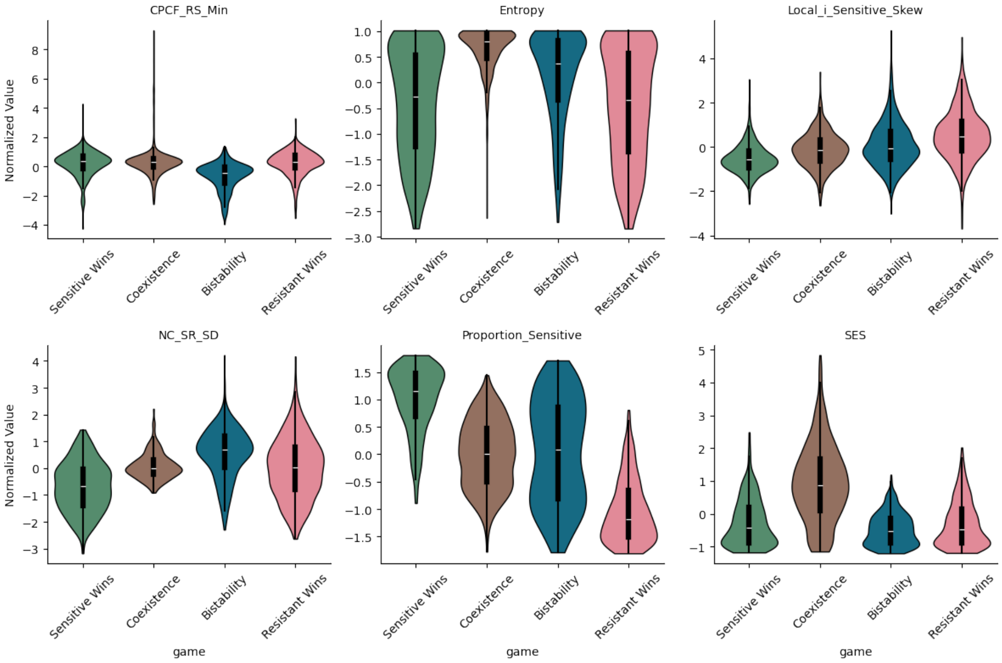


**Fig 7. Feature distributions (normalized) for all features in Main Figure 3.**

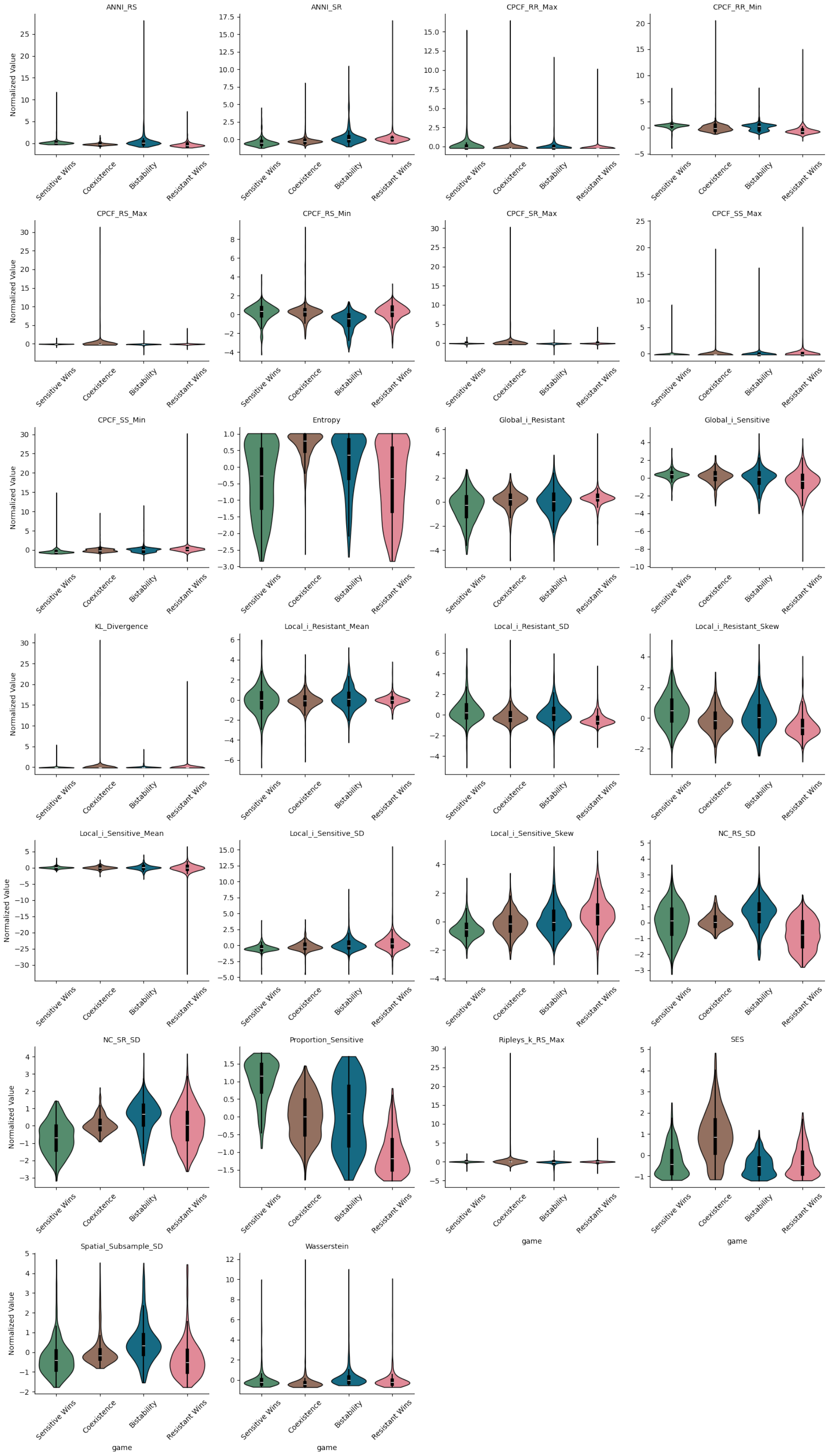


**Fig 8. Feature distributions (normalized) for all non-correlated features.**

#### 5 Mutual information between feature and game

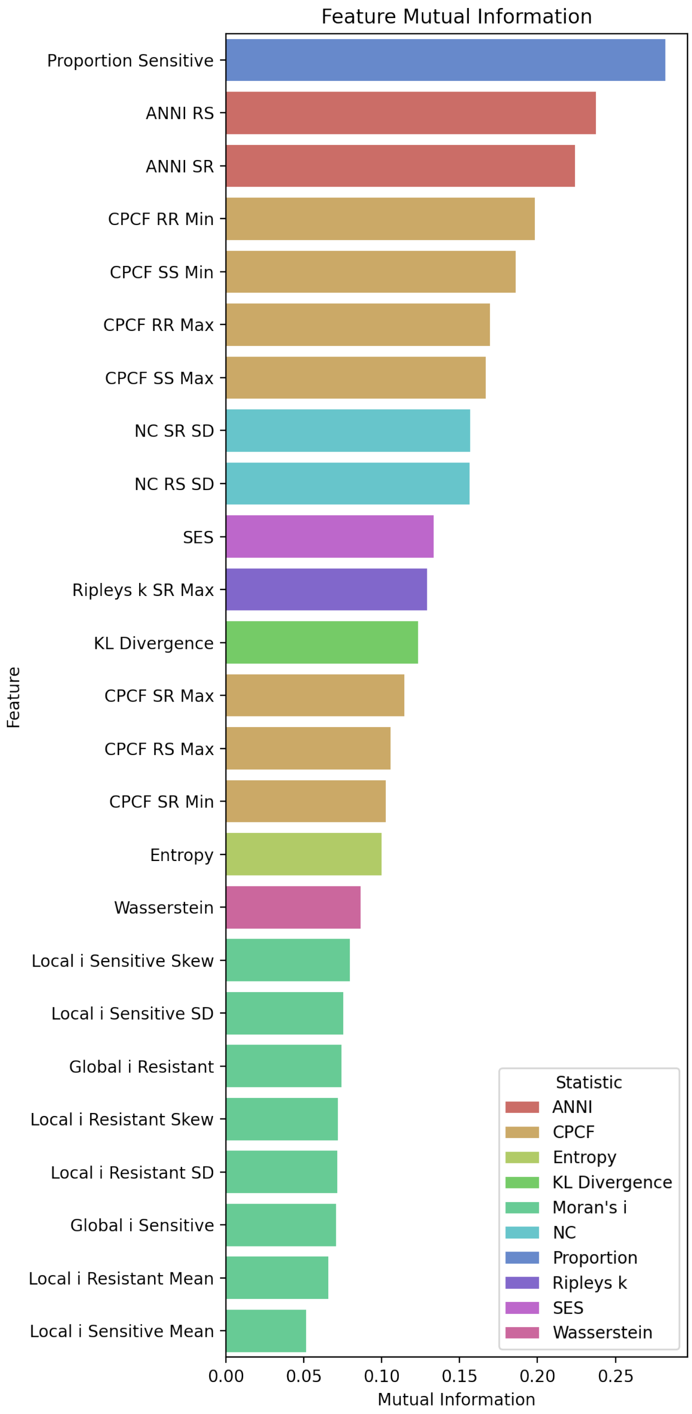


**Fig 9. Mutual information between feature and game, for each feature.**

### 6 Machine learning performance

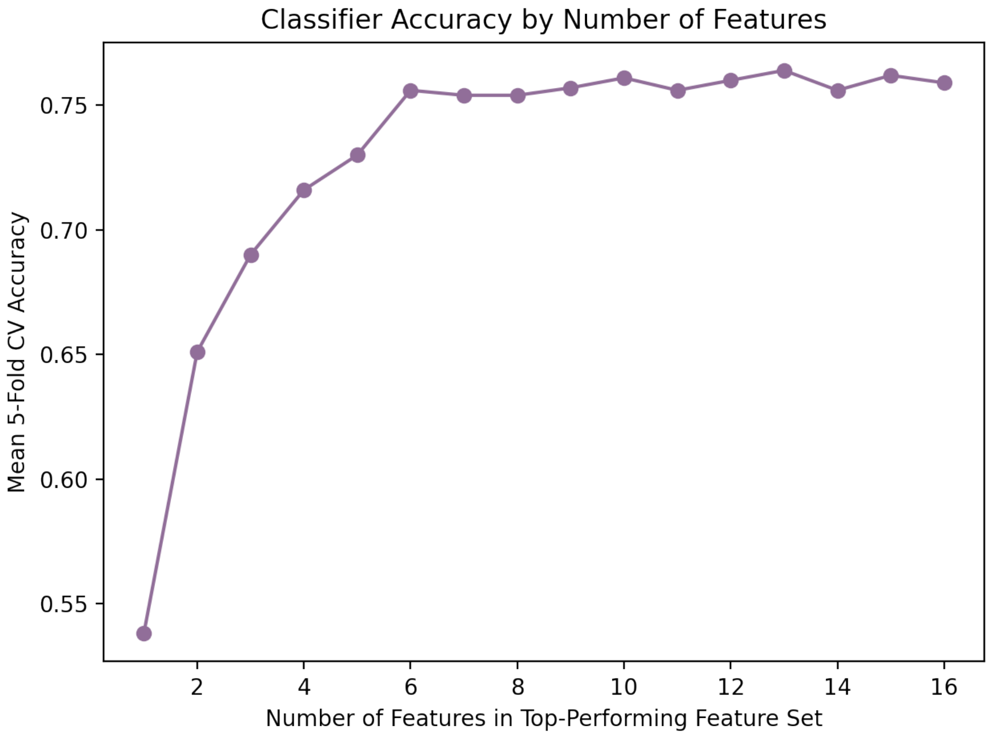


**Fig 10. The 5-fold cross-validation testing accuracy of the highest performing feature set for each feature set size.**

| **Feature** |
|---|
| ANNI RS |
| ANNI SR |
| CPCF RR Max |
| CPCF RR Min |
| CPCF SR Max |
| CPCF SS Min |
| KL Divergence |
| Local i Resistant Mean |
| NC RS SD |
| NC SR SD |
| Proportion Sensitive |
| Ripleys k RS Max |

**Table 3. Top-performing feature set based on top-ten sequential feature selection.** The table is ordered by the order features were included in each subsequent feature set as the features were being selected.

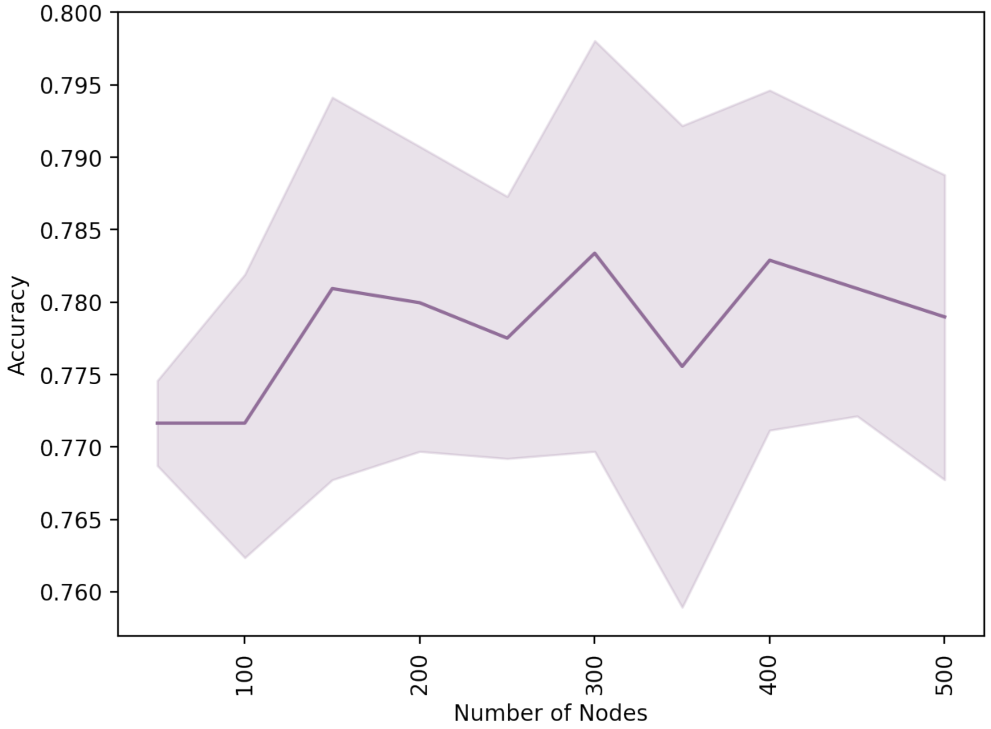


**Fig 11. The 5-fold cross-validation testing accuracy of the model trained on the top 12 features across different number of nodes in the layer.**

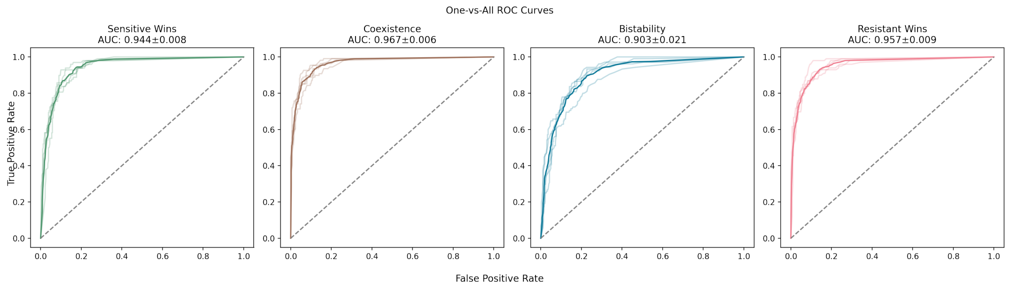


**Fig 12. ROC curves from 5-fold cross-validation runs of the model.**
